# A Familial Alzheimer’s Disease Associated Mutation in Presenilin-1 Mediates Amyloid-Beta Independent Cell Specific Neurodegeneration

**DOI:** 10.1101/2023.07.19.549777

**Authors:** Mahraz Parvand, Joseph JH Liang, Tahereh Bozorgmehr, Dawson Born, Alvaro Luna Cortes, Catharine H. Rankin

## Abstract

Mutations in the presenilin (*PS*) genes are a predominant cause of familial Alzheimer’s disease (fAD). An ortholog of *PS* in the genetic model organism *Caenorhabditis elegans (C. elegans)* is *sel-12*. Mutations in the presenilin genes are commonly thought to lead to fAD by upregulating the expression of amyloid beta (Aβ), however this hypothesis has been challenged by recent evidence. As *C. elegans* lack amyloid beta (Aβ), the goal of this work was to examine Aβ-independent effects of mutations in *sel-12* and *PS1/PS2* on behaviour and sensory neuron morphology across the lifespan in a *C. elegans* model. Olfactory chemotaxis experiments were conducted on *sel-12*(*ok2078*) loss-of-function mutant worms. Adult *sel-12* mutant worms showed significantly lower levels of chemotaxis to odorants compared to wild-type worms throughout their lifespan, and this deficit increased with age. The chemotaxis phenotype in *sel-12* mutant worms is rescued by transgenic over-expression of human wild-type *PS1*, but not the classic fAD-associated variant *PS1_C410Y_*, when expression was driven by either the endogenous *sel-12* promoter (*Psel-12)*, a pan-neuronal promoter (*Primb-1*), or by a promoter whose primary expression was in the sensory neurons responsible for the chemotaxis behavior (*Psra-6*, *Podr-10*). The behavioural phenotype was also rescued by over-expressing an atypical fAD-linked mutation in *PS1* (*PS1_ΔS169_*) that has been reported to leave the Notch pathway intact. An examination of the morphology of polymodal nociceptive (ASH) neurons responsible for the chemotaxis behavior also showed increased neurodegeneration over time in *sel-12* mutant worms that could be rescued by the same transgenes that rescued the behaviour, demonstrating a parallel with the observed behavioral deficits. Thus, we report an Aβ-independent neurodegeneration in *C. elegans* that was rescued by cell specific over-expression of wild-type human presenilin.

## 1. Introduction

There are a variety of disorders that lead to impaired memory and cognition among older adults, of which Alzheimer’s disease (AD) is the most common, accounting for 60% of all dementias [1]. The majority of AD and related dementias are chronic illnesses; they appear in stages, and patients’ symptoms worsen with time [2]. There are two forms of AD: sporadic AD where genetic risk factors are inherited in a non-Mendelian fashion, and familial AD (fAD) where genetic risk factors are autosomal dominant [3]. Familial AD presents earlier than the sixth or seventh life decade and is much less prevalent than AD, however, because fAD offers known genetic causes, this form of the disorder has been the target of much research. Mutations in three genes have been linked to a majority of fAD cases: presenilin 1 (*PS1*), and presenilin 2 (*PS2)* and amyloid precursor protein (*APP*; [4]). Although these genes each alter a large number of cellular processes, multiple studies have focussed on a two common effects: mutations in any one of these genes lead to increased production of amyloid-beta (Aβ) and inflammation [5].

The presenilins have a number of intracellular functions; the most commonly studied is their role as the catalytic subunit of γ-secretase, a protein responsible for the proteolysis of many proteins including APP and Notch [6]. To date, 350 PS1 mutations have been reported in AD patients [7]. A second protein, PS2, has significant homology to PS1 at both gene and protein levels and has also been identified as a cause of fAD [8]. Mutations in PS2 account for the smallest percentage of fAD cases and lead to a later age of onset compared to PS1 and APP mutations [4]. In the clinic, the majority of presenilin mutations identified in patients are missense substitutions that present an autosomal dominant inheritance pattern. As dominant phenotypes typically imply a gain-of—function mechanism, it is believed that pathogenic mutations in PS lead to a toxic gain-of-function phenotype that increases Aß42/Aß40 ratios, a well-known biomarker of AD [9]. However, more recent analysis using mammalian cells has shown that while a vast majority of the studied PS1 mutations do lead to increased Aß42/Aß40 ratios, a majority of these variants significantly decrease production of both Aß42 and Aß40 and have no significant correlation to mean age at onset of fAD for the corresponding mutations [10], casting doubt on the widely accepted assumption that pathogenic PS1 variants lead to fAD by favoring production of Aß42 over Aß40.

In studies of the underlying causes of AD, APP has been a major focus of research. The amyloid hypothesis was formalized for the first time in 1992 when Hardy and Higgins proposed that the build-up of plaques containing Aβ is the predominant cause of AD [11]. Since then, many studies have attempted to determine how amyloid plaques lead to AD and whether elimination of plaques would cure AD. Thus far, research on amyloid plaques has not led to clear answers or to an effective treatment for AD. Although amyloid plaques are considered a hallmark of AD, a build-up of amyloid peptides can also be part of normal aging as Aβ plaques are apparent in 45% of cognitively normal elderly individuals [12]. Because the strongest predictor of both Aβ build up and AD is aging, it may be that other aging-related issues combine with the build-up of amyloid plaques to trigger AD. Since the aggregation of Aβ plaques cannot, on its own, be the sole causal mechanism underlying AD, researchers are exploring the notion that AD is a complex disorder and that multiple pathways other than amyloid plaques may produce the disease characteristics [13].

Because of the known role of presenilins in processing APP, much of the research done on *PS1*/*PS2* is within the context of processing APP. Mouse models containing only *PS1* or *PS2* mutations show an increased proportion of Aβ_42_ but do not exhibit amyloid plaques, thus numerous rodent studies investigating PS1 have used double transgenic mice that have mutations in both *APP* and *PS1* transgenes [14,15]. Plaque formation and cognitive impairment phenotypes manifest much earlier and much more extensively in these double transgenic lines, compared to *APP* transgenic lines without *PS1*/*PS2* mutations or transgenes [14].

Somewhat less studied in rodents are phenotypes in transgenic lines expressing only presenilin mutations, possibly due to lack of complete AD-like pathology. However, recent rodent work has shown that PS1 and PS2 fAD transgenic lines show a plethora of phenotypes, including increased susceptibility to hippocampal damage from kainite induced excitotoxicity or trimethyltin treatment, increased protein oxidation and lipid peroxidation in the mutant brain, impaired hippocampal neurogenesis in adult mice and an age-dependent impairment of spine morphology and synaptic plasticity in hippocampal neurons [15,16]. Partial or complete loss of presenilin by the knock-out of both PS1 and PS2 leads to age-dependent neurodegeneration in the mouse brain [17–19], and in another PS1 knock-in line aging-dependent neurodegeneration and neuronal loss has also been observed [20]. In electrophysiological studies, PS1 fAD mutations were found to alter long-term potentiation (LTP) in the hippocampus [16]. In studies comparing very young transgenic mice over-expressing an fAD mutation in PS1 to counterparts over-expressing wild-type *PS1,* fAD mutants showed enhanced LTP. However, at 8-10 months of age LTP was similar to wild-type mice, and at 13-14 months of age LTP was significantly decreased [16]. Pharmacological blocking studies also showed that the fAD-associated *PS1* mutants altered NMDA receptor-mediated neurotransmission [21]. Behavioural experiments in young mice have also shown that PS1 fAD transgenic mice have behavioural impairments, although the deficits observed are somewhat subtle and inconsistent [16]. Taken together, there is evidence to suggest an alternate hypothesis that loss-of-function mutations in the presenilin genes can elicit AD pathology through one or more molecular pathway(s) independent from that for processing APP [22].

Another proteolytic target for γ-secretase is Notch. Alterations in Notch proteolysis by mutations in PS1 that affect γ-secretase may also be involved in AD pathogenesis. The Notch protein is a type I transmembrane cell surface receptor that plays a role in cell fate decisions of both vertebrates and invertebrates [23,24]. Binding to a member of the Delta, Serrate, or Lag2 (DSL) family of ligands triggers Notch proteolysis, leading to the cleavage of Notch by many of the same secretases that cleave APP [25]. Notch is expressed in neurons at especially high levels in the hippocampus in the adult rodent brain [26], and in some rodent models with presenilin deficiencies, Notch loss-of-function phenotypes are observed [27,28]. Thus, there is evidence to suggest that alterations in Notch can be a contributing factor to AD pathogenesis in the context of PS1. To date the role of Notch signaling in AD pathogenesis remains largely unclear.

In the present study, we used the roundworm *Caenorhabditis elegans (C. elegans)* to investigate the role of PS1 and Notch in neural degeneration and relevant behavioral impairments produced by a mutation in *sel-12*, an orthologue of *PS1.* SEL-12 shares approximately 50% amino acid identity with human PS1, among which include many AD-associated residues. As *C. elegans* express two genes orthologous to human Notch receptors, *glp-1* and *lin-12*, questions regarding Notch signaling in the context of fAD can be interrogated in this model system as well. There are a number of advantages to using *C. elegans* for studying neurological degenerative diseases including its short life span (14-18 days), a sequenced genome, and defined nervous system with 302 identified neurons that are well characterized with a mapped connectome [29]. Basic cellular neurobiological functions are well conserved, with *C. elegans* having cellular and biochemical processes similar to those in mammals. *C. elegans* shows both genetic and functional similarity with orthologs of many of the mammalian neurotransmitters (e.g., dopamine, serotonin, GABA, glutamate, acetylcholine, and their receptor subtypes) and many classes of ion channels (e.g., sodium, calcium, and potassium channels) [30]. Several known components of vesicle release were first identified in *C. elegans* (*i.e* UNC-18), and their mammalian orthologs discovered later (MUNC-18) [31]. Moreover, the first known function of presenilins was the role of PS1 in cleaving Notch based on the observation that mutations in *sel-12* produced an egg-laying deficit known to be caused by impaired Notch signaling [32]. Lastly, because the *C. elegans* ortholog of *APP, apl-1*, does not generate a product orthologous to Aβ, the nematode model offers a unique opportunity for studying functions of presenilins in an Aβ-free biological system.

Here we studied whether mutations in *sel-12* altered chemotaxis and sensory neuron morphology across aging, and whether deficits could be rescued by over-expressing variants of human *PS1*. To do this we first tested olfactory chemotaxis across development and during aging in *sel-12* mutant worms, and in worms over-expressing either human wild-type *PS1* or one of the _two *PS1* mutations:_ *PS1_C410Y_,* a classical fAD mutation that impairs both Ab and Notch processing, [33,34], and a novel mutation, *PS1_ΔS169,_* which is believed to impair Ab but leave Notch processing intact [35]. In *sel-12* mutants and worms over-expressing the *PS1* transgenes, we also assessed whether there was neurodegeneration of the ASH chemosensory neurons that are responsible for sensing the odorant used in our chemotaxis assay. We found that in *sel-12* mutant worms, there was more degeneration of the ASH sensory neurons with increasing age and this degeneration could be rescued by both nervous system and cell-specific expression of wild-type *PS1* and *PS1_ΔS169_*, but not *_PS1C410Y_*. These results show an Ab-independent, cell specific role of presenilin in neuronal degeneration.

## 2. Materials and Methods

### 2.1 Generation of transgenic lines and strain maintenance

*C. elegans* Bristol wild-type (N2), *sel-12(ok2078)* mutant (RB1672), and the transgenic nuIs[*osm-10::GFP*+*lin-15(+)*] (HA3) strains were provided by the Caenorhabditis Genetics Center (CGC), and the transgenic strain HA1712 (*lin-12 (null); Phsp-16.2::glp-1RNAi*) was provided graciously by Dr. Anne Hart of Brown University. Worms were cultured on *Escherichia coli (E. coli*) seeded nematode growth medium (NGM). All experiments were conducted in 6cm Petri plates that were filled with 10 mL NGM agar a maximum of two weeks prior to use. Worms were stored in a 20℃ incubator and all experiments were conducted in a room with controlled humidity (40 ± 5 % RH) and temperature (20 ± <u>1℃</u>).

Transgenic overexpression lines were constructed by microinjection of DNA plasmids into the germ line of young adult *sel-12* mutant worms at a concentration of 10 ng/μl along with a co-injection GFP marker [36]. This DNA mixture was injected into the distal arm of the worm’s gonad, which contains a central cytoplasm core that is shared by germ cell nuclei [36]. Worms were injected under an inverted Zeiss DIC microscope equipped with a 40X Nomarski objective. The plasmid pBY140 containing the wild-type *PS1* coding region expressed by the *sel-12* promoter was provided by Dr. Ralf Baumeister (Albert-Ludwig University in Freiburg/Breisgau, Germany). To generate pan-neuronal, ASH-specific and AW-specific over-expression constructs, promoter regions of rimb-1 (2.6kb fragment, 734bp upstream of start codon, from pSF11[ptag-168p::nCre]), sra-6 (1.8kb fragment, 158bp upstream of start codon) and odr-10 (986bp fragment, 28bp upstream of start codon), respectively were used to replace the sel-12 promoter region in the original pBY140 via restriction cloning. Lines with >50% GFP transmission were selected for testing. Because DNA microinjection creates an extra-chromosomal array, there is inherent variability of expression levels because there were probably different copy numbers of plasmids incorporated into the array and there was also the possibility of different cells having different genotypes (mosaic expression) [37]. To overcome this, three independent lines for each construct were tested [38].

**List of strains generated and tested:** Please refer to Table 1 of supplemental material.

### 2.2 Chemotaxis Assays

To ensure that all animals tested were the same developmental age, adult worms (n=20-30) were placed in a droplet of 4μl of bleach solution (1:1 ratio of 100% bleach and 1M NaCl). The worms’ bodies disintegrated but their eggs survived. All eggs hatched at roughly the same time, and larvae were allowed to grow for three days on agar seeded with *E. coli* in a 20℃ incubator. On day three, 72-hour-old worms were collected using 700μl of liquid M9 buffer into 1.5mL centrifuge tubes. To remove OP50 *E. coli* from the bodies of the worms, worms were centrifuged for a minute at 1000rcf and the supernatant was removed. Another 700ul of M9 buffer was added to the tubes, and the worms were centrifuged again (done 3 times). After the third time, the supernatant was removed and the pellet of the worms remained in the tube.

The chemotaxis procedure used was adapted from Margie *et al.* [39]. Eight NGM Petri plates (6cm diameter) were used per strain (4 control, 4 test) and each was divided into four equal quadrants. In the center of each plate a circle of 1cm diameter (Fig. 1A) was drawn on the bottom of the plate and lines were drawn to mark the quadrants. For test plates, each quadrant was marked as control or test. For control plates, all 4 quadrants were labelled as control. The octanol solution was prepared by mixing of a 1:1 ratio of 100% octanol and 1M-sodium azide (NaN_3_) that was used to immobilize worms. The diacetyl solution was prepared by diluting it to 0.5% diacetyl using 99.5μl of M9 and 5μl of diacetyl. A 1:1 ratio of this 0.5% diacetyl solution was then added to NaN_3_. Control solution was prepared by mixing equal volumes of M9 and NaN_3_. Washed worms (50-100) were pipetted into the circle at the center of the plates. Then 2μl of the volatile odorants octanol or diacetyl were placed on spots located at the centre of the test quadrants, and 2μl of the control solution was placed on spots located at the centre of the control quadrants (Fig. 1A).

**Figure 1.**
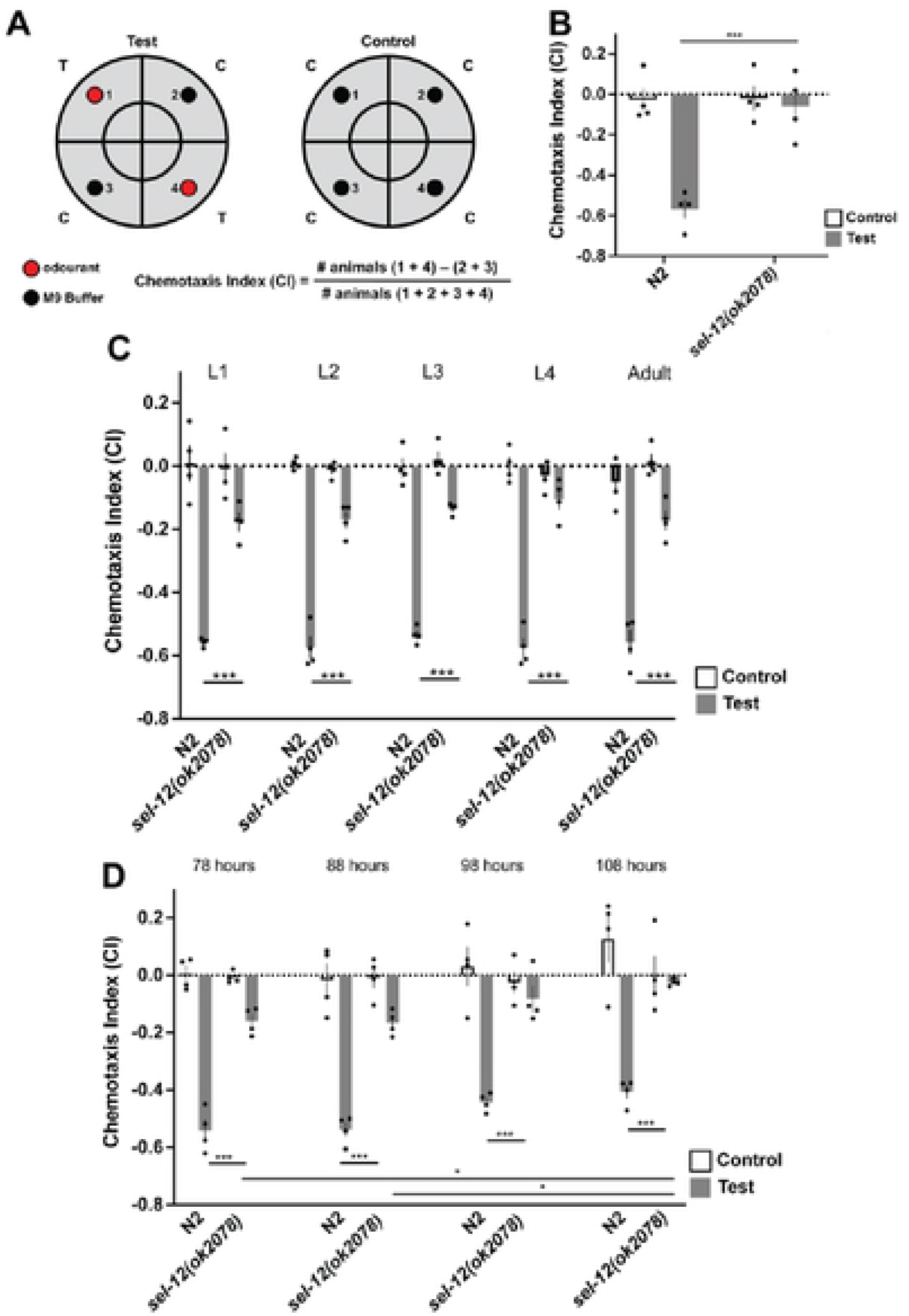
Worms with mutations in orthologs of *PS1* show progressive chemosensory impairments. **A)** Chemotaxis assay-worms are placed in the centre circle and allowed 1 hour to navigate plate. Anesthetic is included in buffer and odorant spots, so worms remain at the quadrant of choice. Worms in each quadrant are counted and CI calculated as shown. **B)** at 72 hours of age, *sel-12* mutants display impaired octanol avoidance behavior. **C)** *sel-12* chemotaxis deficits are present from hatch. **D)** *sel-12* chemotaxis deficits become more severe with age. At 78 hours of age worms are egg laying adults. They continue to lay eggs for 3-4 days. By 7 days of age worms are moving into old age. Each bar represents an average from 4 independent plates (n=50-100 worms per plate). Independent plate CI are shown as circles. White bars represent control plates and black bars represent test plates. Error bars reflect the standard error of the mean.*** p<0.001, *p<0.05

Plate lids were kept open for approximately 15 minutes until the odorants soaked into the agar and the surface of the agar appeared dry. Worms were then left to move around the plates undisturbed for one hour in a 20°C room. Plates were then transferred into a 4°C refrigerator for two hours to immobilize worms before counting. Prior to counting plates were relabeled and a key generated, plates were removed from the refrigerator for counting one at a time and the number of worms in each quadrant was counted by a researcher blind to the condition/strain/the study objectives. After counting the blind was broken and a chemotaxis index (CI) was calculated (Fig. 1A).

### 2.3 Use of 5’-fluorodeoxyuridine (FUdR) in aging studies

Because *sel-12* mutant worms have an egg-laying deficit, eggs accumulate inside the gonad and the progeny hatch inside the adult killing it at a relatively young age (Levitan et al., 1995). This makes studies using older worms difficult. To stop the hatching of eggs inside adults so the adults lived to an older age, we used 5’-fluorodeoxyuridine (FUdR). FUdR is an inhibitor of DNA synthesis that is used in *C. elegans* lifespan studies to preserve a synchronized aging population [40]. Two days prior to adding *E. coli* to NGM plates, 250μl of 100μM FUdR was spread on plates. Two days after the addition of *E. coli*, Larval stage 4 (L4, approx. 50 hours post hatch) animals were transferred to the NGM plates with FUdr. For aging studies, adult worms were tested at 10 hour intervals from 78 to 108 hours (young adult to senior worms). In experiments where FUdR was used all genotypes were treated with FUdR, including the wild-type control.

### 2.4 Locomotion Assay

Behavioral experiments on worms of different ages were used to determine speed and the locomotory ability of worms on plates to ensure they were able to chemotax to the test spots in the time given. The multi-worm tracker (MWT) is an automated machine vision system that records features of worm behavior for many worms simultaneously; for this assay crawling speed was measured [41]. To assess locomotion, approximately 50 worms of each age were handpicked onto a plate without FUdR or *E. coli* (n=3 plates), placed on the MWT, and the lid of each plate was lifted briefly to provide an air-puff stimulus to arouse worms so they were moving and could be recognized by the MWT. Worms were tracked for 250 seconds.

### 2.5 Knock-down of both Notch receptors *via* heat shock and developmental stage synchronization

Because worms carrying null mutations in both *glp-1* and *lin-12* have a lethal phenotype, we studied the effect of double loss-of-function of both Notch receptors by the adulthood induction of *glp-1* RNAi knockdown using a heat-shock promoter (*hsp-16.2*) in a *lin-12(n941)* null mutant background. Animals with a heat shock promoter driving *glp-1* RNAi were raised at 15°C, the permissive temperature. During heat-shock experiments, all strains used, including wild-type, were moved to 33°C, the restrictive temperature, for 2 hours as described in Singh *et al*. [42] After this, animals were allowed to recover for 3 hours in a 20°C incubator prior to chemotaxis experiments that were conducted in a 20℃ behavior room. Expression levels of *hsp-16.2* persist even up to 50 hours after heat shock [43], suggesting that after 3 hours of recovery there would still be sufficient knockdown of *glp-1*. Note that Notch mutant worms have developmental delays compared to wild-type. To test worms at similar stages in development, wild-type and *lin-12/glp-1* deficient worms were closely observed every 4 hours to determine when these strains reached young adulthood (defined as containing at least 4 eggs [44]). Compared to wild-type worms, worms with deficiencies in both Notch receptors had a 6-hour lag in development (reached young adults at 68 hours in 20°C).

### 2.6 ASH Neuron Imaging

The single pair of ASH sensory neurons in the head of the worm are the primary chemosensory receptors for detecting octanol. For imaging we used strains expressing *Posm-10::GFP* that filled the ASH neuron cell bodies and processes with GFP and imaged both wild-type and *sel-12* mutant worms over time (78, 98, and 108 hour old worms). For all extrachromosomal rescue strains, the line that had the closest chemotaxis index to wild-type was chosen for imaging. The researcher doing the imaging was blind to the strains being imaged. Worms were placed on 2% agar pads on sterile glass microscope slides in 15uL of 50mM NaN_3_ for immobilization. Worms were given approximately one minute to become immobile prior to being covered with a 1.5mm thick coverslip. Images were obtained using a Leica SP8 white light confocal microscope. To excite GFP, a 488nm wavelength laser was used and the emitted light was collected by passing through a 510-550nm bandpass filter. Depending on the thickness of the worm and the brightness of its GFP, optical sections were collected at various intervals using a 63X oil immersion lens and summed into a single Z projected image. Neurons were quantified using the Fiji: ImageJ Software.

Independent groups of one hundred worms were imaged per strain at 78, 98 and 108 hours of age for a total of 300 images per strain. During analysis, ASH neuron morphology was classified into three categories: normal, gapped/missing arm, and bleb. If gaps and blebs were both present, the neuron would be categorized as blebs.

For DiI dye-filling experiments, animals were collected and incubated in 400μL 1.25μM DiI in the dark for 30 minutes. After incubation, worms were transferred back onto OP-50 seeded NGM plates for 30 minutes then collected for imaging.

### 2.7 Statistical Analysis

Data are reported as means ± standard error of the mean (SEM) of three or four independent plates with 50-100 worms on each for each strain tested. Because expression levels from extrachromosomal transgenes can be variable, we analyzed data for 3 independent lines of each transgene. For the sake of clarity, we report analysis for our collapsed data. The presenilin rescue strains are reported as the average of four plates for each of three extrachromosomal lines (total of 12 plates). Each rescue line was analyzed independently and then as averages of the three extrachromosomal rescue lines for each strain. Data analyses were conducted using the GraphPad Prism software, SPSS and python (pingouin 0.5.1). Data were checked for normal distribution and one-tailed independent T tests were conducted for pair-wise comparisons. In experiments with three or more groups, a one-way ANOVA was conducted with a Tukey’s Honest Significant Difference test with statistical significance set at p=0.05 for post-hoc analyses. The percentage of normal and abnormal ASH morphology of each strain was compared to other strains using aggregated Pearson’s chi-squared test. P values of less than 0.05 were considered significant. Data visualization was done with GraphPad Prism, Python (with Plotly 5.7.0 graphing library) and Adobe Illustrator.

## 3. Results

### 3.1 *C. elegans* with a *sel-12* mutation displayed olfactory impairments shortly after hatching that increased over time

In *C. elegans*, there are three members of the Presenilin family: *hop-1, sel-12, and spe-4* [38,45,46]. *spe-4* is only distantly related to human presenilins and is expressed only in the male germline [45], and thus was not investigated in this work. Although both *hop-1* and *sel-12* have homology to human *PS1* and are widely expressed throughout the worms, including in muscles and neurons [32], *sel-12* has higher sequence homology to *PS1* than *hop-1* (50% vs 31%). For this reason, we focused our work on worms expressing a putatively null *sel-12(ok2078)* deletion allele.

To assess the olfactory abilities of wild-type and *sel-12* mutant worms across the lifespan we examined them across development (Fig. 1C). *C. elegans* have 4 larval stages (lasting from ∼12-24 hrs in length) and we tested the aversive chemotaxic response to octanol at each stage, L1-L4, as well as at 78 hours old (reproductive young adult worms). Compared to wild-type worms, *sel-12* mutants had significant chemotaxis index deficits from L1 that were also seen in L2, L3, L4, and adult worms [F (9, 30) = 65.60, p <0.001] (Fig. 1C).

We also tested whether the olfactory deficits progressed with advanced age by testing worms ranging in age from reproductive adult to old age at 10 hour intervals (78-108 hours; Fig. 1D). The multivariate analysis of covariance showed a significant group effect indicating there was a significant difference in CI scores between groups ([F(7, 24) = 60.11, p<0.001]). Post-hoc analyses showed that although there was a mild but non-significant pattern of decrease in chemotaxis index over time between 78 hours old and 108s hour old in wild-type worms (p=0.06), the decrease in chemotaxis index was much more rapid and was significant in *sel-12* mutant worms across age (Fig. 1D; p=0.05). Overall, wild-type and *sel-12* mutant worms had chemotaxis index decrements from 78 to 108-hours old of 15% and 81%, respectively.

### 3.2 Motor ability is not a confounding variable in *sel-12(ok2078)* mutant chemotaxis

It is possible that the egg-laying phenotype of the *sel-12* mutants could lead to bloating and impair the ability to locomote well enough to successfully chemotax within the allotted time. We investigated this possibility by assessing locomotor speed of wild-type and *sel-12* worms at 68hr, 78hr, 88hr, 98hr and 108hrs of age on the MWT, in a neutral *E. coli* (OP-50) seeded environment without any odourants. Although at 68 hours old *sel-12* mutant worms appeared slower than wild-type by visual observation, via MWT analyses average forward speed during the entire tracking session was not significantly different (Supp. Fig. 1A; p=0.08). In comparing worms aged from 68 hours to 108 hours old, wild-type worms’ forward speed peaked at 88 hours and slowed down at 108 hours old. In contrast, the speed of *sel-12* mutant worms did not increase from 68 hours to 88 hours old and was significantly slower at 88-hour (t(4)=5.89, p=0.002) and 98-hour old (t(4)=6.10, p=0.002) compared to wild-type worms (Supp. Fig. 1A). Despite the observation that *sel-12* mutant worms move more slowly than wild-type worms, they were able to navigate around the entire petri plate within approximately 5 minutes (Supp. Fig. 1B), demonstrating that they could exhibit chemotaxis behavior in the 1-hour test session if they were able to detect the olfactory cue. Thus, despite a marked reduction in locomotion speed in comparison to their wild-type counterparts, *sel-12* mutants maintained sufficient locomotor ability to explore and navigate to all areas of the plate environment in the given time.

### 3.3 Expression of wild-type *sel-12* and wild-type *PS1* rescued chemotaxis deficit when driven by the endogenous *sel-12* promoter

Using extrachromosomal arrays we over-expressed wild-type *sel-12* with its endogenous promoter (*Psel-12::sel-*12) in the *sel-*12 mutant and showed that it rescued octanol chemotaxis behavior (Fig. 2A; [F(2,17) = 2.65, p<0.001]). Over-expressing human *PS1* driven by the *sel-12* promoter (*Psel-12::PS1_WT_*) also rescued the chemotaxis deficit. Chemotaxis tests were also conducted on wild-type worms, *sel-12* mutant worms, and rescue lines over-expressing *Psel-12::PS1_WT_* in the *sel-12* mutant background at 88, 98, and 108 hours of age (Supp. Fig. 2). For each age, *Psel-12::PS1_WT_* rescue lines restored chemotaxis deficits in *sel-12(2078)* mutant worms and had CI values significantly different from *sel-12* mutant worms (At 88 hours [F(2,17) = 67.98, p<0.001], at 98 hours [F(2,17) = 33.92, p<0.001], and at 108 hours [F(2,17) = 64.73, p<0.001]). This finding was consistent with a previous study testing functional conservation between *sel-12* and PS1 in which over-expressing human wild-type PS1 using the endogenous *sel-12* promoter in a *sel-12* mutant rescued mutant phenotypes [32]. Because *sel-12* is expressed in both neuronal and non-neuronal cells in the worm, we next tested whether the octanol chemotaxis deficit could be rescued by expressing PS1 in just the nervous system.

**Figure 2.**
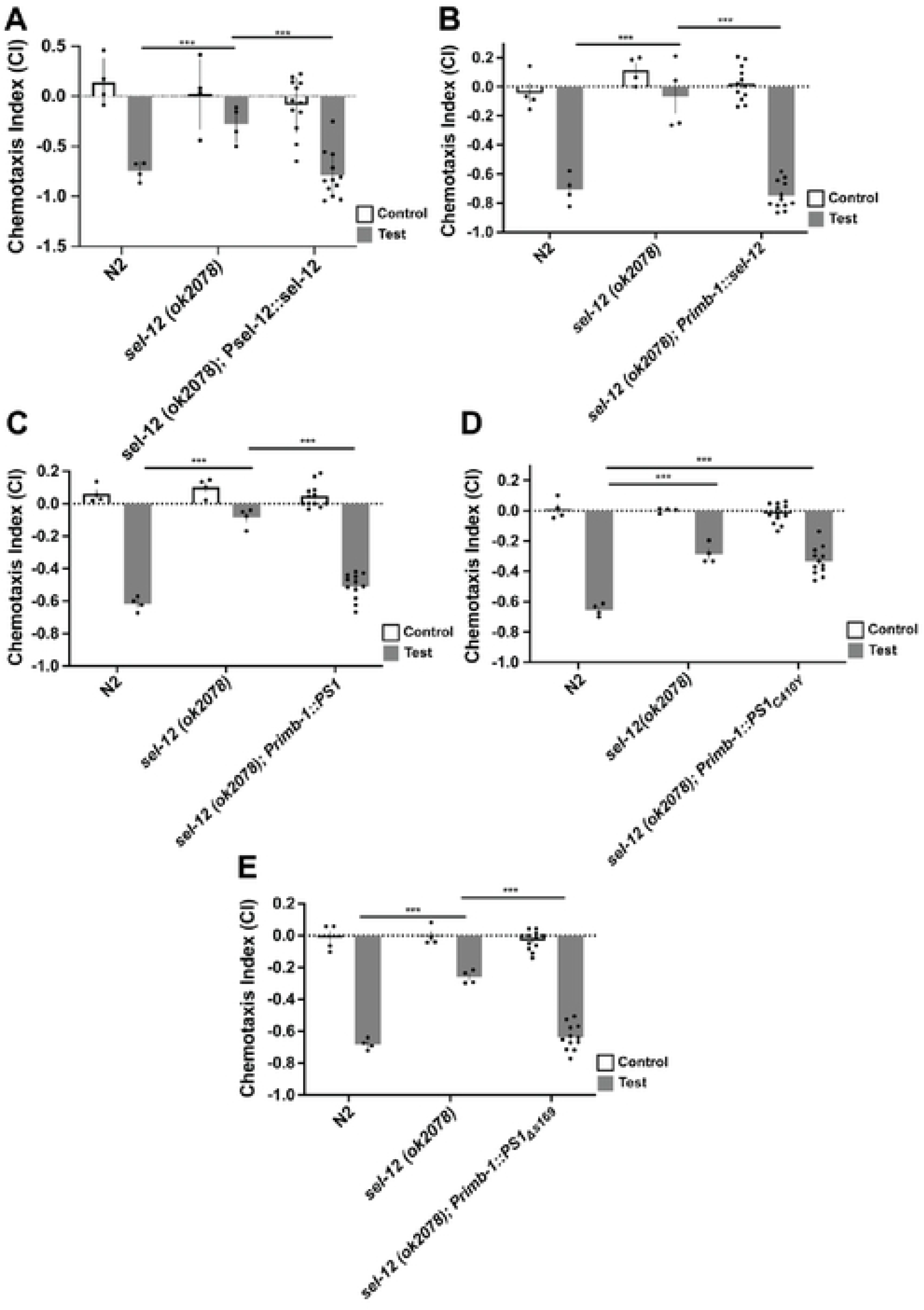
*Psel-12-driven* expression of human wildtype PS1 and pan­neuronal expression of wildtype genes and *PS1_Δs169_* rescued octanol chemotaxis deficits in *sel-12* mutant worms. A) *Psel-12::sel-12* rescued chemotaxis deficits of *sel-12* mutants. B) *Psel-12::PS1* rescued chemotaxis deficits of sel-12 across age. B-D are done with 72 hr old animals. C) *Ptag-168::sel-12* rescued chemotaxis deficits. D) *Ptag-168::PS1* rescued chemotaxis deficits. E) *Ptag-168::_PS1C410Y_* did <u>not</u> rescue chemotaxis deficits. F) *Ptag­168::PS1_Δs169_* rescued chemotaxis deficits. Octanol was used for test (aversive odorant) and M9 for controls (odorless buffer). Each bar represents an average of 4 independent plates (n=50-100 worms per plate). Rescue bars are collapsed averages of 3 independent rescue lines generated. Independent plate CI are shown as circles. White bars represent control plates and black bars represent test plates. Error bars reflect the standard error of the mean. *** p<0.001.

### 3.4 Nervous system expression of wild-type *sel-12*, wild-type *PS1*, and *PS1_Δs169_,* but not *PS1_C410Y,_* rescued chemotaxis deficits in *sel-12* mutant worms

In these experiments we used *rimb-1 (*formerly *tag-168)*, which encodes a pan-neuronally expressed presynaptic active-zone protein [47], to over-express extrachromosomal arrays of wild-type *sel-12,* wild-type *PS1*, and *PS1_Δs169_,* and *PS1_C410Y,_* pan-neuronally and tested chemotaxis. *Primb-1::sel-12* rescued the chemotaxis deficits in *sel-12* mutant worms (Fig. 2B; [F(2,17) = 41.68, p<0.001]; wild-type vs *sel-12* rescue p=0.32, *sel-12* mutant vs *sel-12* rescue p<0.001). Pan-neuronal over-expression of wildtype human PS1 (*Primb-1::PS1_WT_)* also rescued chemotaxis deficits in *sel-12* mutant worms (Fig. 2C; [F(2,17) = 68.23, p<0.001]; wild-type vs *PS1* rescue p=0.44, *sel-12* mutant vs. *PS1* rescue p<0.001).

Interestingly, over-expressing the fAD-linked *PS1* mutation C410Y in the nervous system of *sel-12* mutant worms (*Primb-1::PS1_C410Y_*) did not rescue chemotaxis deficits in *sel-12* mutant worms (Fig. 2D; [F(2,17) = 27.67, p<0.001]; wild-type vs. *PS1_C410Y_* rescue p<0.001, *sel-12* mutant vs. *PS1_C410Y_* rescue p=0.59). These findings are also consistent with those of Levitan et al. (1996), who showed that fAD-associated PS1 variants were unable to rescue the *sel-12* abnormal vulva and egg-laying phenotypes. Finally, we tested whether the novel fAD-linked mutation, *PS1_Δs169_*, had an effect on chemotaxis. It was previously reported that *PS1_Δs169_* impacts APP processing and amyloid generation without affecting Notch1 cleavage and Notch signalling as the egg-laying phenotype in *sel-12* worms could be rescued by *PS1_Δs169_* [35]. Interestingly, in this study over-expressing *Primb-1::PS1_Δs169_*rescued chemotaxis deficits in *sel-12* mutant worms (Fig. 2E; [F(2,17) = 52.24, p<0.001]; wild-type vs. *PS1_Δs169_* rescue p=0.31, *sel-12* mutant vs. *PS1_Δs169_* rescue p<0.001).

In *sel-12* mutant animals an attractive chemotaxis response to diacetyl was also impaired. Using the same strains, we saw that over-expressing *sel-12*, *PS1_WT_* and *PS1_Δs169_*pan-neuronally also rescued impaired diacetyl chemotaxis, while expressing *PS1_C410Y_*did not (Supp. Fig. 3A-E).

### 3.5 ASH neuron-specific expression of wild-type *sel-12*, wild-type *PS1,* and *PS1 _Δs169,_* but not *PS1_C410Y_*, rescued chemotaxis deficits in *sel-12* mutant worms

In *C. elegans*, the ASH sensory neurons are polymodal nociceptors that detect a number of noxious and potentially toxic stimuli including octanol [48]. The avoidance response to octanol is mediated by ASH activation [48]. To determine whether presenilin plays a role in chemotaxis by acting specifically in these sensory neurons, we over-expressed *sel-12, PS1_WT_*, *PS1_C410Y_,* and *PS1_Δs169_* in *sel-12(ok2078)* mutants driven by promoter region of *sra-6*, a gene strongly expressed in the ASH neurons, and weakly expressed in the ASI neurons and PVQ interneurons [49]. Over-expressing *Psra-6::sel-12* rescued chemotaxis deficits in *sel-12* mutant worms (Fig. 3A; [F(2,17) = 45.26, p<0.001]; wild-type vs *sel-12* rescue p=0.21, *sel-12* mutant vs. *PS1* rescue p<0.001).

**Figure 3.**
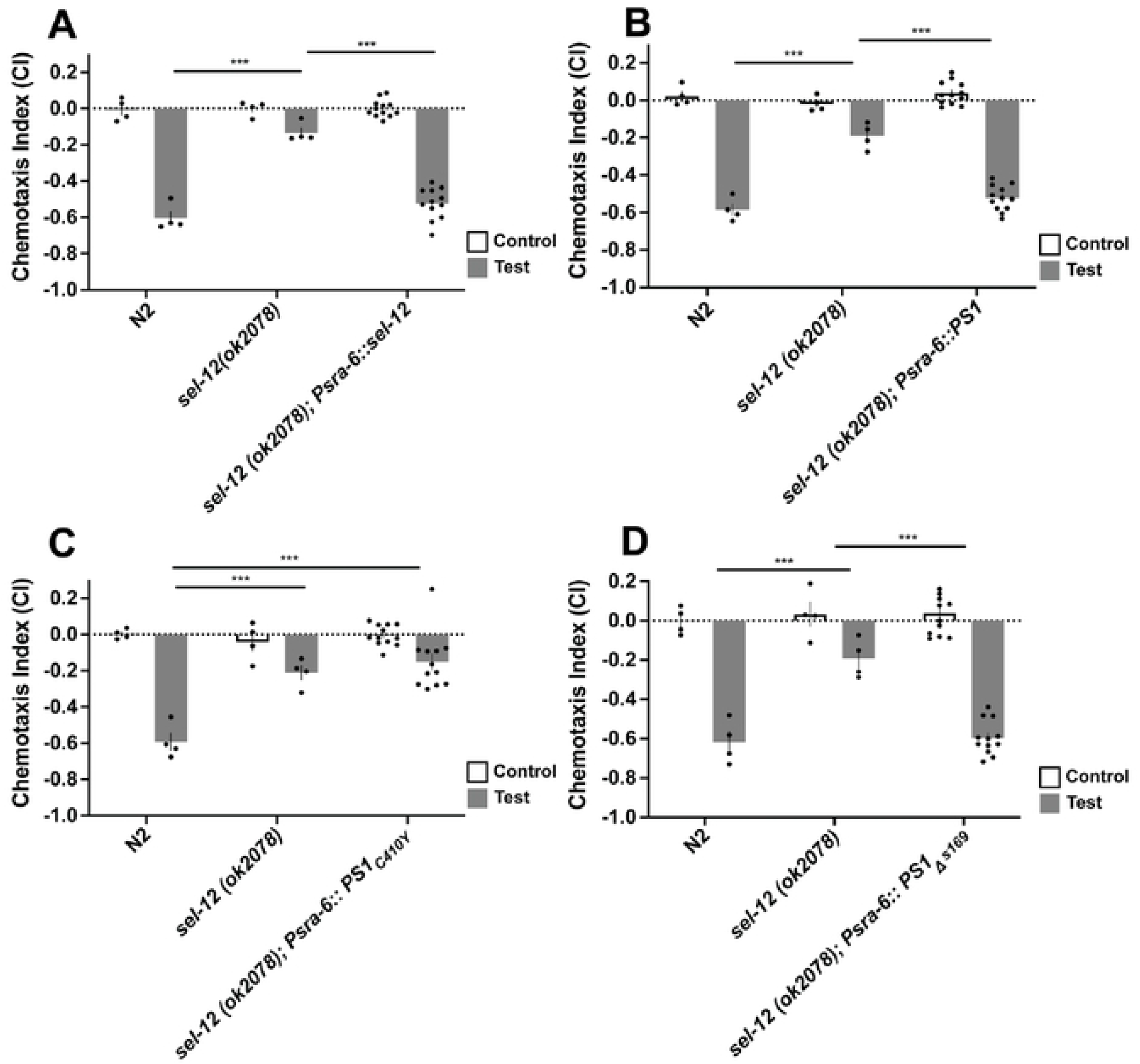
ASH-neuron specific expression of wildtype genes and *PS1_Δs169_* rescued octanol chemotaxis deficits in *sel-12* mutant worms. A) *Psra6:: sel-12* rescued chemotax.is deficits. B) *Psra6::PS1* rescued chemotaxis deficits. C) *Psra6::PS1_C410Y_* did <u>not</u> rescue chemotaxis deficits. D) *Psra6:: PS1_Δs169_* rescued chemotaxis deficits. Octanol was used for test (aversive odorant) and M9 for controls (odorless buffer). Each bar represents an average from 4 independent plates (n=50-100 worms per plate). Rescue bars are collapsed averages of 3 independent rescue lines generated. Independent plate CI are shown as circles. White bars represent control plates and black bars represent test plates. Error bars reflect the standard error of the mean. *** p<0.001

Next, we over-expressed wild-type human *PS1* under the control of the *sra-6* promoter (Fig. 3B), and found that it rescued the chemotaxis impairments observed in *sel-12* mutant worms ([F(2,17) = 44.29, p<0.001]; wild-type vs *PS1_WT_* rescue p=0.28, *sel-12* mutant vs. *PS1* rescue p<0.001). On the other hand, when we expressed the canonical fAD *PS1* mutation C410Y in the ASH neurons (*Psra-6::PS1_C410Y_*; Fig. 3C), we found that it did not rescue chemotaxis deficits in *sel-12* mutant worms ([F(2,17) = 16.68, p<0.001]; wild-type vs. *PS1_C410Y_* rescue p<0.001, *sel-12* mutant vs. *PS1_C410Y_* rescue p=0.72). Finally, in contrast to *Psra-6: PS1_C410Y_* over-expression, *Psra-6::PS1_Δs169_* over-expression did restore chemotaxis deficits in *sel-12* mutant worms (Fig. 3D; [F(2,17) = 30.69, p<0.001]; wild-type vs. *PS1_Δs169_* rescue p=0.92, *sel-12* mutant vs. *PS1_Δs169_* rescue p<0.001). This data aligns with previous studies that show that there is functional conservation between human PS1 and its nematode ortholog. The loss-of-function mutations of both PS1 and its nematode ortholog appear to impair cell function in a cell autonomous manner as over-expressing these genes in a single pair of sensory neurons rescued the *sel-12* chemotaxis deficit.

### 3.6 Wild-type *PS1* rescue lines in the ASH neuron had a cell-specific effect on chemotaxis

Although wild-type *PS1* in ASH was sufficient to rescue octanol chemotaxis, it is possible that the rescue was not cell-autonomous, and that expression of *PS1_WT_*anywhere in the nervous system would rescue the phenotype. To rule out this hypothesis, we expressed wild-type *PS1* driven by the *odr-10* promoter that is expressed in another pair of chemosensory neurons, the AWA neurons, which are responsible for chemotaxis towards the attractant diacetyl. Consistent with our ASH-octanol avoidance findings, worms with the same mutation in *sel-12* are also deficient in appetitive chemotaxis towards diacetyl and they show the same pattern of chemotaxis deficits and rescues for PS1_WT_ and PS1 mutations as was observed for aversive chemotaxis away from octanol (Supp. Fig. 4A-C). We tested chemotaxis away from octanol and towards diacetyl in wild-type worms, *sel-12* worms and worms with wild-type *PS1* expressed in either the AWA neurons or the ASH neurons of *sel-12* mutant worms.

We then tested whether AWA-specific over-expression would rescue ASH-specific phenotypes, and whether ASH-specific over-expression would rescue AWA-specifc phenotyes. Over-expression of wild-type *PS1* in the AWA neurons rescued the chemotaxis defect towards _diacetyl_ (Fig. 4B; [F (2,17) = 100.21, p<0.001]; wild-type vs *PS1_WT_* rescue p=0.76)_, but did not_ rescue chemotaxis impairment away from octanol (Fig. 4A; [F(2,17)= 137.30, p<0.001]; wild-type vs *PS1_WT_* rescue p<0.001). Additionally, over-expressing wild-type *PS1* in the ASH neurons did not rescue *sel-12* mutant animals’ impaired chemotaxis towards diacetyl but did rescue the chemotaxis away from octanol (Fig. 4C; [F (2,17) = 54.00, p<0.001]; wild-type vs PS1 rescue p<0.001).

**Figure 4.**
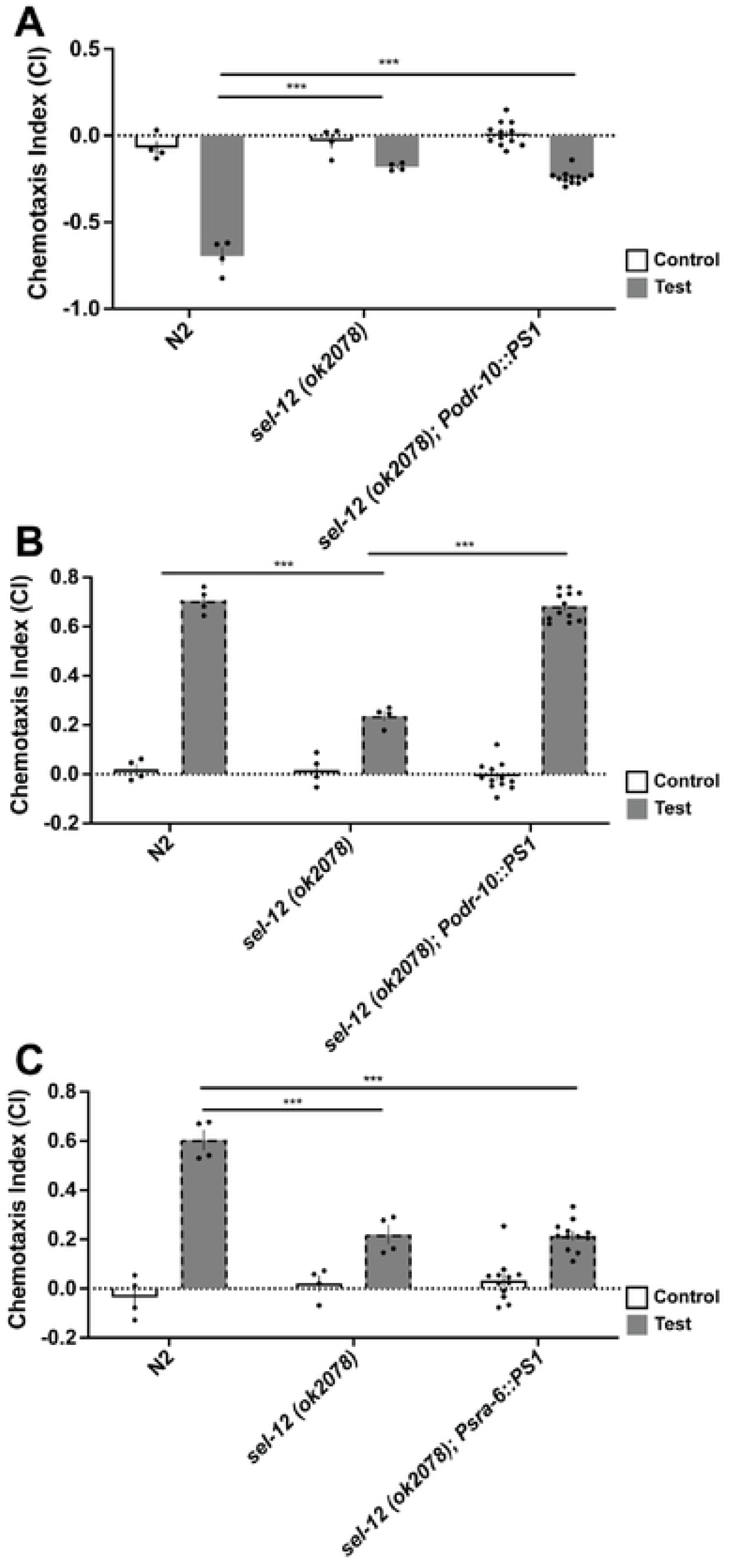
Wildtype *PS1* over-expression in the ASH neuron had a cell-specific effect on rescuing chemotaxis in *sel-12* mutant background. Wildtype *PSI* was expressed in AWA neurons by odr-10 promoter (responsible for diacetyl) and ASH neurons by sra-6 promoter (responsible for octanol) cell-specifically. **A)** *Podr-10::PS1* did not rescue chemotaxis deficits in response to octanol. **B)** *Podr-10::PS1* rescued chemotaxis deficits in response to diacetyl. **C)** *Psra6::PS1* did not rescue chemotaxis deficits in response to diacetyl. Non-dotted bars denote chemotaxis assay with done octanol, while dotted bars denote a chemotaxis assays done with diacetyl. Each bar represents an average from 4 independent plates (n=50-100 worms per plate). Rescue bars are collapsed averages of 3 independent rescue lines generated. Independent plate CI are shown as circles. White bars represent control plates and black bars represent test plates. Error bars reflect the standard error of the mean. *** p<0.001

Thus, expression of *PS1_WT_* in the ASH neurons rescued chemotaxis deficits in *sel-12* mutant worms for the octanol odorant, but not for the diacetyl odorant sensed by different sensory neurons, the AWA neurons. At the same time expression of *PS1_WT_* in the AWA sensory neurons did not rescue chemotaxis to octanol, but did rescue chemotaxis to diacetyl. Thus, via double-dissociation we show that the roles of *sel-12* in mediating the chemotaxis deficits are cell specific, and cell-autonomous.

### 3.7 *C. elegans* with mutations in Notch receptors do not show increased chemotaxis deficits over time

Contrasting chemotaxis data from animals with the two fAD mutations, the classic mutation *_PS1C410Y_* that alters both Aβ and Notch cleavage, and *_PS1Δs169_* mutation that affects Aβ cleavage but leaves Notch intact, suggested an important role for Notch cleavage by *sel-12* in normal chemotaxis. Like APP, Notch receptors are cleaved and activated by the presenilins of γ-secretase. In earlier research Singh *et al.* reported that in *C. elegans* the two Notch receptors, *lin-12* and *glp-1*, regulate chemosensory avoidance behavior in response to octanol [42]. They found that a knockdown of *glp-1* in a *lin-12* null background led to behavioral impairments in response to octanol. We used this same approach to investigate whether the octanol chemosensory deficit observed in Notch mutants showed the same characteristics as the *sel-12/PS1* mutation. As single mutations of either *glp-1* or *lin-12* did not show chemotaxis impairments in our experiments (data not shown), we used the transgenic strain created by Singh *et al.* in which *glp-1* was knocked down using heat-shock promotor driven RNAi in adult *lin-12(n941)* mutants, and tested octanol chemotaxis of this strain as worms aged to determine whether these deficits increased in the same way as did chemotaxis in the *sel-12* mutant worms.

Both *sel-12* mutant worms and *lin-12/glp-1* mutant worms had significantly lower CIs compared to wild-type worms (p <0.001; Fig. 5). Compared to age-matched wild-type worms, *C. elegans* with *lin-12*(n941)*/glp-1* reduction-of-function did not show increasingly impaired octanol chemotaxis over time (p=1.00). This is in contrast with wild-type (N2) and *sel-12* mutant worms that demonstrated an increasing chemotaxis deficits over time (p<0.001). These findings suggest that Notch associated chemotaxis deficits do not fully phenocopy all of the characteristics of mutations in *sel-12*.

**Figure 5.**
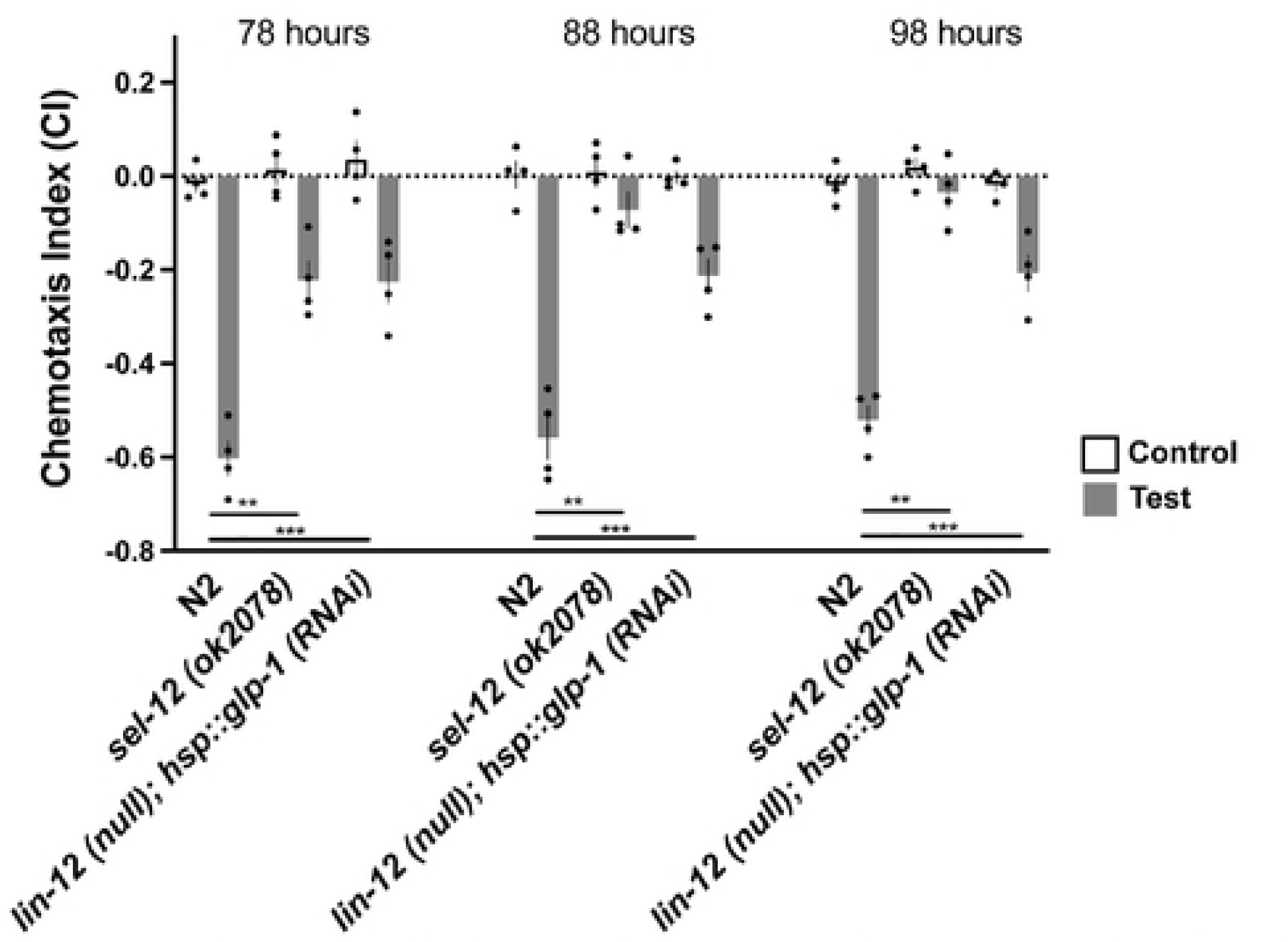
Worms with RNAi knockdown of *glp-1* in a *lin-12* null background showed chemotaxis deficits that did not progress over time. Octanol was used for test (aversive odorant) and M9 for controls (odorless buffer). Each bar represents an average from 4 independent plates (n=50-100 worms per plate). Independent plate CI are shown as circles. White bars represent control plates and black bars represent test plates. Error bars reflect the standard error of the mean. *** p<0.001

### 3.8 *sel-12* mutant *C. elegans* have ASH neuron morphological abnormalities that are rescued by wild-type *PS1* and *PS1_Δs169_*, but not *PS1_C410Y_*

The chemotaxis experiments showed that *sel-12* mutant worms had chemotaxis deficits that increased as worms aged. What causes the chemotaxis deficits? Since neuronal death is observed in AD we hypothesized that there might be neuronal degeneration occurring in *sel-12* worms. This led us to investigate whether the chemotaxis behavioral phenotype reflected morphological changes or neurodegeneration of the sensory neurons. To address whether the loss-of-function mutation in *sel-12* altered the morphology of the ASH neurons, we crossed the *sel-12(ok2078)* mutant into a strain of worms expressing cytosolic GFP in the ASH neurons (osm-10::GFP), and using this resultant strain as the background generated transgenic lines over-expressing *PS1_WT_*, *PS1_Δs169_* and *PS1_C410Y_*. Using a confocal microscope we imaged the ASH neurons in separate groups of 100 wild-type, *sel-12* mutant, *Psra-6::PS1_WT_, Psra-6::PS1_Δs169, and_ Psra-6::PS1_C410Y_* worms at 78, 98, and 108 hours of age. A number of different phenotypes were observed that were categorized into “normal”, “gapped”, or “blebs” (Fig. 6A). ASH neurons can also be visualized via a DiI dye-filling protocol [50]. When we looked at dye-filling in our GFP strains, the GFP co-localized with DiI dye in the ASH neurons, suggesting that the blebbing and gap phenotypes observed are indeed indicative of degenerated processes and not due to non-uniform distribution of GFP (Fig. 6B). Within the same strain, there appeared to be no significant differences in the total number of abnormal ASH neurons across different ages (Fig. 6C; p>0.05 between all ages within the same strains). A chi-square test showed that *sel-12* worms had significantly fewer normal ASH neurons compared to wild-type worms across all time points, *X^2^* (1, N=200) = 16.33, p<0.01. However, as worms aged there appeared to be a progression from a “gapped arm” state to a more severe “blebbed” state in degenerating neurons – and this progression appeared more prominent in *sel-12* mutants and PS1_C410Y_ over-expression animals. Animals with ASH neuron-specific expression of wild-type *PS1* and *PS1_Δs169_*, had significantly more normal neurons than *sel-12* animals and were not significantly different from wild-type, however animals with *PS1_C410Y_* were not significantly different from *sel-12* animals and had significantly fewer normal neurons than wild-type animals (Fig. 6C). As a proxy measure for the health of the neurons the area of GFP in each neuron was quantified. ASH neurons in that wild-type worms had significantly greater area of GFP compared to *sel-12* mutant and *Psra-6::PS1_C410Y_* worms at all time points; in *sel-12* mutant animals the decrease in GFP area across age is particularly noticeable (Fig. 6D). These data support the hypothesis that in *sel-12* mutants the ASH neurons were degenerating as worms aged.

**Fig 6.**
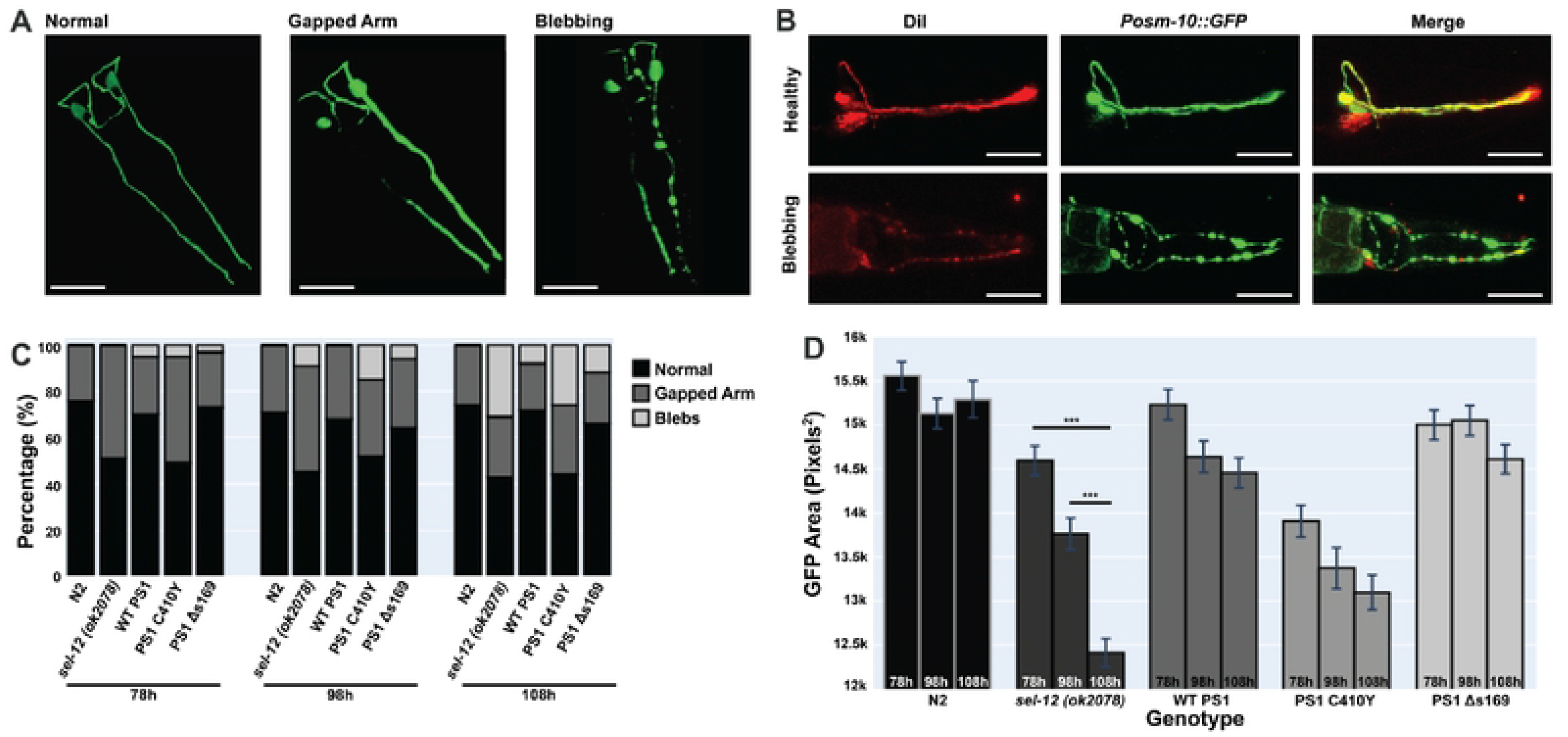
*sel-12* mutant worms have impaired ASH sensory neuron morphology that was rescued by ASH­ specific expression of wild-type and *PS1_Δs169_*. **A)** Categories of ASH sensory neuron morphology. ASH neurons have been categorized as healthy with normal morphology, having gaps in the processes or have circular blebs. Both gaps and blebs are indicative of neurodegeneration. Scale bar = 30µm **B)** In healthy ASH neurons, dye-fill with Dil overlaps with *Posm-10::GFP.* In degenerating ASH neurons, there is little or no overlap between Dil dye-fill and the fluorescent marker. Scale bar= 30µm. **C)** ASH neuron morphology of five strains (n=100 per strain per time point) overtime (78, 98, and 108 hour old worms). Different worms were used for each time point. ASH neurons of wildtype, *sel-12* mutant, ASH-specific wildtype *PS1* rescues, ASH-specific *PS1_Δsl69_* rescues, and ASH-specific *PS1_C410Y_* rescues were imaged and categorized as either normal or abnormal (gapped arm or blebs). *X2* (1, N=200) = 16.33, p<0.01. **D)** ASH GFP area quantification in pixels^2^. *** p<0.001.

## 4. Discussion

Our studies expand on previous evidence supporting the hypothesis of an Aβ-independent role of human *PS1* in neural degeneration by studying *sel-12*, the *C. elegans* orthologue of *PS1, in vivo*. We demonstrated that in an Aβ-absent model, C410Y mutations in *PS1* lead to AD-like pathogenesis.

We observed that worms with *sel-12* mutations had chemotaxis deficits from shortly after hatching, and that these deficits increased with age (Fig. 1D). The ASH neurons that are responsible for detection of the octanol odorant showed morphological abnormalities indicative of neurodegeneration and degenerated more rapidly in *sel-12* mutant worms compared to wild-type worms (Fig. 6). This progressive neurodegeneration paralleled the decline of the chemotaxis indices of *sel-12* mutant worms over time. Human PS1 and *sel-12* have considerable conservation of function, such that the over-expression of human WT PS1 rescued *sel-12* mutant phenotypes [32]. Here we found that the over-expression of either wild-type *PS1* and *PS1_Δs169_*_, but not *PS1*_*_C410Y_*, in the ASH neurons rescued both chemotaxis deficits and ASH morphological abnormalities in *sel-12* mutant worms (Figs. 2 & 6). As *PS1_Δs169_* is hypothesized not to affect Notch signaling while *PS1_C410Y_* does, we investigated whether the interaction between *PS1* and Notch played a role in these chemotaxis deficits. Previously, Singh et al. [42] reported that worms with decreased expression of both of the two Notch receptors *lin-12* and *glp-1* showed octanol chemotaxis deficits, suggesting that Notch is involved in mediating chemotaxis behaviours. By using their approach and knocking down *glp-1* with RNAi in a *lin-12* mutant strain, we confirmed the importance of Notch for the chemotaxis phenotype. However, we also found that the chemotaxis deficits induced by impairments in Notch signaling did not increase as animals aged (Fig. 4). This suggests that although Notch plays an important role in octanol chemotaxis, *PS1* may be functioning in chemotaxis in an additional, Notch-independent manner. Both wild-type *PS1* and *PS1_Δs169_* rescued neuron morphology, suggesting that human and worm presenilins and Notch have conserved functions in post-mitotic neurons.

Because *sel-12* is strongly expressed in all *C. elegans* neurons, these data raise the question of whether the entire nervous system degenerates as a result of *sel-12* mutations. Our behavioural data suggest that, in addition to the ASH neurons, the AWA sensory neurons may also be degenerating. Previous work on *sel-12* worms examined the AIY interneurons and the mechanosensory neurons (ALM and PLM). Wittenburg *et al.* [51] showed that *C. elegans* with mutations in *sel-12* have defects in temperature memory as a result of a loss of function in cholinergic interneurons from the time of hatching [51]. They also showed that these mutant worms displayed abnormal morphologies in the AIY interneurons, and that this phenotype was rescued by AIY-specific wild-type *sel-12* and wild-type *PS1* expression [51]. Sarasija *et al.* [52] reported that mutations in *sel-12* lead to metabolic defects of the mitochondria, resulting in oxidative stress and neurodegeneration in the ALM and PLM neurons responsible for responding to mechanical stimuli [52]. Reducing calcium levels in the mitochondria in *sel-12* mutant worms prevented neurodegeneration in mechanosensory neurons [52]. Therefore, it is possible that mutations in *sel-12* impact a number of neurons. Our locomotion data suggests that not all neurons are affected in mature *sel-12* mutant worms as they were still able to move about on the agar surface, indicating that the motor neurons continue to drive movement even in the oldest worms we tested (Supp. Fig. 1A-B). *C. elegans* provides a unique system in which the role of presenilin in olfactory dysfunction and neuron death can be investigated using single cell rescues. As presenilins are substrates for different caspase proteins, the interaction between *PS1* and caspases can be further explored. This use of single cell resolution phenotyping allows the characterization of even subtle and pleotropic phenotypes, providing potential novel therapeutic opportunities.

While findings from our experiments are consistent with other observations in the literature that demonstrate mechanistic loss-of-function phenotypes of PS1 function, our experiments do not reconcile this with the autosomal dominant nature of pathogenic PS mutations in fAD. In our experiments transgenes were over-expressed as extrachromosomal arrays leading to variability between the transgenic lines. We do not know the copy number of the transgenes expressed by these arrays (it could be a few copies or several hundred copies), and the expression level would likely have been much higher than what is found *in vivo.* This suggests to us that overexpressed levels of wildtype *sel-12* and human PS1 do not cause any deficits in chemotaxis in *C. elegans*. Newer techniques such as the Mos10 mediated Single Copy Insertion (MosSCI) [53] and the CRISPR-Cas9 system can be used in *C. elegans* to generate single or low number copy rescues [54]. In MosSCI, the mobilization of a transposon creates DNA double strand breaks in the non-coding region, which gets repaired by using the extrachromosomal template to copy DNA into a chromosomal site [53]. In CRISPR-Cas9, a spacer sequence is transcribed into an RNA sequence that can then locate matching DNA sequences. When this is done, the Cas9 enzyme can either activate or deactivate gene expression [54]. Both approaches would allow a single copy of PS1 to replace mutant *sel-12* in worms and might produce more consistent effects. Additionally, this approach can make testing phenotypes in a heterozygote possible and further resolve the paradox between the mechanistic loss-of-function hypothesis of PS and the autosomal dominant mode nature of pathogenic mutations of PS. Interestingly, in mice studies inactivation of a single PS1 locus is not sufficient to cause neurodegeneration while having one copy of a pathogenic variant is [55,56] – it is hypothesized, therefore, that the resulting mutant protein is antimorphic and in addition to being loss-of-function can act in a dominant-negative manner to inhibit the activities of normal PS produced from the remaining alleles to cause the dominant inheritance pattern of PS mutations. This hypothesis has yet to be fully explored.

This work together with previous studies [52,62] demonstrate that neurodegeneration of neurons in *C. elegans* with *sel-12* mutations can occur in the absence of Aβ deposits. This offers an opportunity to study the role presenilins may play in neurodegeneration, independent of Aβ. Data from Sarasija et al. [52] suggested that the *sel-12* neural degeneration was due to mitochondrial dysfunction that was Notch independent, however their tests for Notch looked only at single mutations in *glp-1* and *lin-12* and they did not test a double knock-down to test for redundancy. Our data suggests that although Notch contributes to the chemotaxis deficit and neural degeneration phenotypes, double loss or knock-down of Notch receptors shows that the age-progressive phenomenon may be Notch-independent. In addition, our data with the new pathogenic PS1 variant that has a functional Notch processing domain, *PS1_Δs169_*, suggests that PS1’s role in fAD pathogenesis possibly extends beyond just the Notch processing pathway. Studies interrogating other molecular pathways PS1 is involved will uncover possible other pathways of PS1 that may contribute to fAD.

Taken together, the results we obtained can best be explained by two possibilities: one, that the fAD mutation has a lethal effect in neurons leading to cell death, or two, that presenilins play an important role in cell maintenance or survival and that classic fAD mutations interfere with that function and without it neurons die. Thus, a pathogenic mutation either kills the cell, or stops an essential function required for the cell to survive. This offers interesting research questions to pursue in the future. Our observation of cell-specific rescue shows that the effect of the mutation is cell specific and so is not a circuit issue, but an issue at the level of individual neurons. *C. elegans* offers the unique opportunity to study the functions of human disease genes and disease-causing mutations at the level of single neurons in living, behaving animals. This knowledge may lead to novel approaches to studying AD and developing new approaches to diagnosis and treatment of this debilitating disorder.

## Acknowledgements

The authors would like to thank the Caenorhabditis Genetics Centre (CGC) for providing the wild-type and *sel-12* mutant strains, Dr. Ann Hart for providing the Notch mutant strains, Dr. Ralf Baumeister for providing the wild-type *sel-12, PS1*, and *_PS1C410Y_* _strains, and Dr. Weihong Song_ for providing the *PS1_Δs169_* plasmid.

This research was supported by a grant from the Canadian Institutes of Health Research #PJT 165947 to CHR, MP was supported by the Ecole Polytechnique Commemorative Award from the Canadian Federation of University Women (CFUW), JL was supported by an NSERC PGS-D.

## Author Contributions

MP and CR designed the studies and MP participated in all experiments. TB piloted and designed the chemotaxis assay MP made and tested the *Psel-12::PS1, Psel-12::PS1C410Y* and *Psel-12::PS1_Δs169_* rescue strains. DB conducted pan-neuronal chemotaxis experiments. JL conducted neuronal imaging. ALC designed all plasmids used in this project. The first draft of the manuscript was written by MP and CR and then rewritten by JL. All authors read and approved the final manuscript.

